# DNA replication in early mammalian embryos is patterned, predisposing lamina-associated regions to fragility

**DOI:** 10.1101/2023.12.25.573304

**Authors:** Shuangyi Xu, Ning Wang, Michael V. Zuccaro, Jeannine Gerhardt, Timour Baslan, Amnon Koren, Dieter Egli

## Abstract

DNA replication in differentiated cells follows a defined program, but when and how it is established during mammalian development is not known. Here we show using single-cell sequencing, that both bovine and mouse cleavage stage embryos progress through S-phase in a defined pattern. Late replicating regions are associated with the nuclear lamina from the first cell cycle after fertilization, and contain few active origins, and few but long genes. Chromosome breaks, which form spontaneously in bovine embryos at sites concordant with human embryos, preferentially locate to late replicating regions. In mice, late replicating regions show enhanced fragility due to a sparsity of dormant origins that can be activated under conditions of replication stress. This pattern predisposes regions with long neuronal genes to fragility and genetic change prior to segregation of soma and germ line. Our studies show that the formation of early and late replicating regions is among the first layers of epigenetic regulation established on the mammalian genome after fertilization.

## Introduction

DNA replication is an essential process for cell division and development, but basic principles are largely uncharacterized in the early mammalian embryo. It was recently shown that DNA replication stress defined by low replication fork speed, replication fork stalling, and a requirement for G2 DNA synthesis, is prevalent in both mouse and human embryos. Particularly in human embryos, replication stress is associated with fork collapse and chromosome breakage. As a result, human embryos frequently incur replication-dependent DNA damage and aneuploidies (*1*). Abnormal chromosome content and DNA damage acquired post fertilization impair the developmental potential of the embryo and are thus relevant for our understanding of developmental failure. Aneuploidies acquired after fertilization are also commonly seen in porcine, rhesus macaques and bovine embryos (*2-5*), while in mice spontaneous aneuploidies are uncommon (*6*). Thus, genome instability appears to be the norm in mammals, with humans and mice at opposite ends of the spectrum for both developmental potential of a fertilized egg and for the incidence of chromosomal abnormalities. Furthermore, viable genetic change during early cell divisions may result in developmental defects in the fetus and disease in the newborn, such as through error-prone fork recovery pathways and chromosomal rearrangements. This has added relevance, as DNA replication stress in human embryos was shown to primarily affect regions containing long genes implicated in neurodevelopmental disorders. The molecular determinants underlying this pattern of chromosome breakage and aneuploidy are unknown.

Replication timing, a temporal order of DNA replication, is thought to play an important role in shaping the genomes of multicellular organisms, with late replicating regions experiencing higher mutation rates and fragility (*7*). In somatic cells, common fragile sites (CFS) are late replicating regions, which are prone to break upon induction of replication stress (*8*). In the early mammalian embryo, replication fork speed is physiologically slow even without added aphidicolin (*1, 9*). Spontaneous DNA breakage occurs in the embryo at sites concordant with those in somatic cells (*1*), raising the question whether DNA replication is also patterned in the mammalian embryo. In frog, fly and fish embryos prior to the midblastula transition, DNA replication is very rapid and near random, with an organized replication program emerging gradually (*10-12*). Such randomness might also be expected in mammalian embryos, as key epigenetic properties of chromatin affecting DNA replication patterns are only being established: chromatin architecture, which is linked to DNA replication patterns (*13*), solidifies during embryonic genome activation (*14*), alongside major epigenetic changes in DNA methylation and histone modifications. However, S-phase in mammalian embryos proceed much slower, taking hours instead of minutes as in lower vertebrates, and the cell cycle in mammalian embryos requires between 15-24 hours to complete instead of less than an hour for zebrafish and frogs. Therefore, whether DNA replication patterns in mammalian embryos have a defined program like in more differentiated mammalian cells cannot be inferred and has not been determined.

While in somatic cells transcription-replication conflicts contribute to replication stress, genome instability in early preimplantation-stage embryos occurs independent of transcription (*1*). Possible determinants of the patterns of fragility may be limiting origin density and late DNA replication, where the consequences of low fork speed and of DNA replication fork stalling and fork collapse may be greatest.

Here we used mouse and bovine embryos to determine the DNA replication timing during cleavage development. We chose zygote and cleavage stage embryos for analysis as it is the developmental point associated with chromosome breakage and might thus provide insight into why the genome of some mammalian embryos shows a specific pattern of fragility. We find that late replicating regions emerge already in the first cell cycle, and a replication timing program is evident at the 2-cell stage and 4-cell stage in mice, as well as in early bovine cleavage stage embryos prior to embryonic genome activation (EGA). Long genes expressed primarily in terminally differentiated cells such as neurons, neuronal gene clusters, as well as long intergenic regions replicate late in the cell cycle. Late replicating regions show nuclear lamina association, low origin density and increased fragility. Thus, the early establishment of DNA replication timing in the mammalian embryo predisposes specific regions to genetic change in the soma and the germ line.

## Results

### Mouse preimplantation embryos show patterned progression of DNA replication

To determine the replication timing profile of mammalian embryos, we analyzed mouse zygotes, 2-cell stage and 4-cell stage cleavage stage embryos, as well as parthenogenetically activated embryos containing only a maternal genome. Cleavage stage embryos were dissociated, and the genome of individual cells was amplified and sequenced (**Fig 1A**). The dataset included in the analysis encompassed individual nuclei from 80 zygotes, and 219 mouse blastomeres harvested at the 2-4 cell stage. Recently developed methods to measure DNA replication timing of single cells were used, based on read frequency mapped to the respective reference genome (*15*). MetaphaseII oocytes were used as reference to even copy numbers across the genome. Individual cells/nuclei are displayed in the order according to the percent genome replicated (**Fig. 1B**). To define early and late replicating regions, replication timing profiles were summarized by counting the number of cells with replicated versus un-replicated DNA at each genomic bin. Here we define late replication as regions where less than 50% of the samples are replicated at each genomic bin. DNA replication timing patterns were beginning to emerge from the zygote stage: maternal and paternal nuclei isolated from fertilized zygotes or single nuclei from parthenogenically activated zygotes showed concordant regions of late DNA replication with blastomeres (e.g. at the Vmn2r region). Theamplitude of the differences between early and late replicating regions at the cleavage stage was greater than at the zygote stage (**Fig.1B**). This suggests that DNA replication at the 1-cell stage has greater stochasticity than later stages. From the 2-cell stage, individual cells/nuclei followed a highly similar pattern of DNA replication, apparent as congruent areas of replicated or unreplicated DNA (**Fig. 1B**). Blastomeres showed an ordered progression from unreplicated to replicated across the genome (**Fig. 1B**, **Fig. S1A**, **S1B**.) Aggregate replication profiles, indicated as ‘summary’, demonstrate distinct early and late replicating regions throughout the genome (**Fig 1B, Fig. S1**).

**Figure 1.**
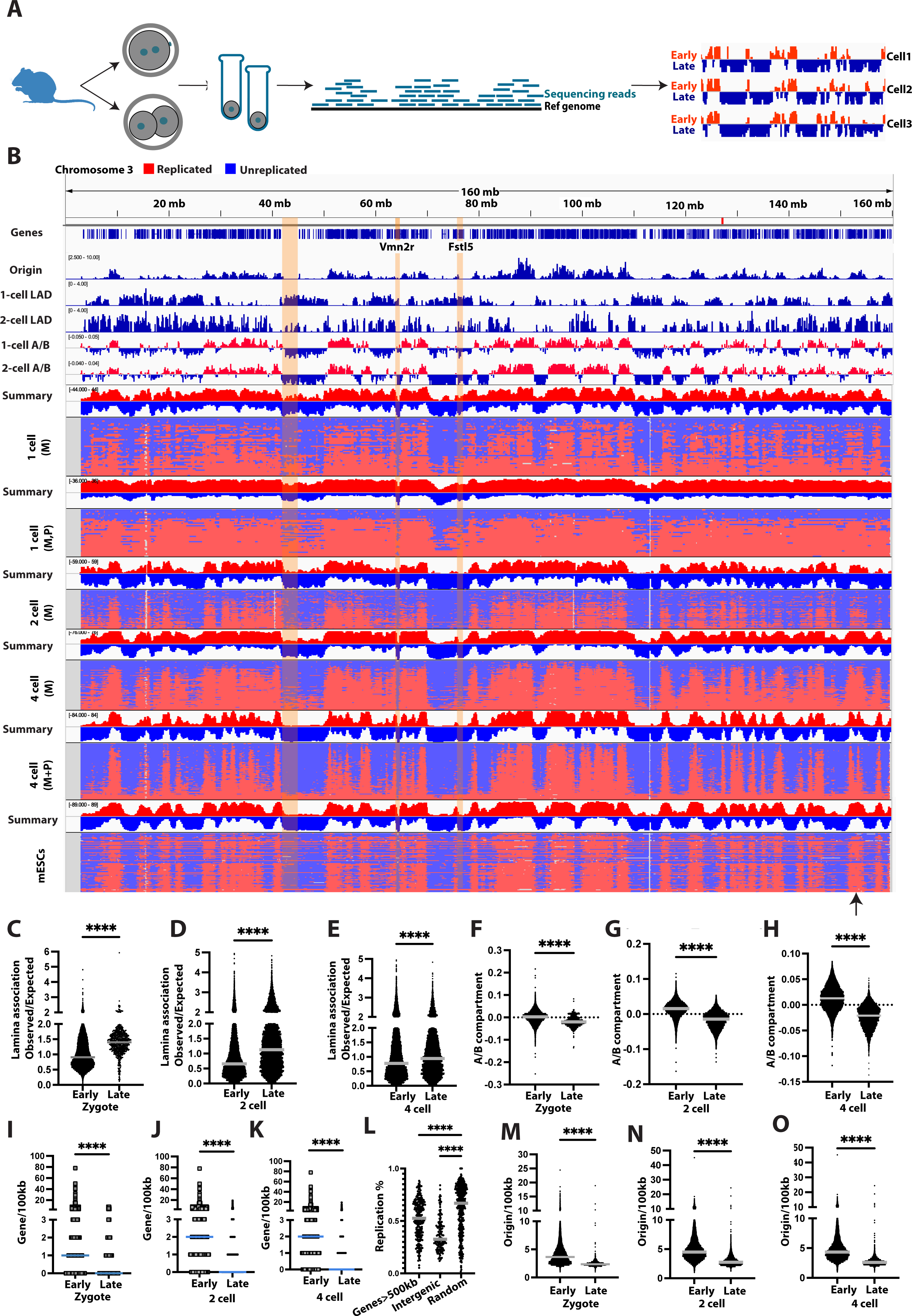
DNA replication timing in mouse embryos is patterned, correlating with gene and origin density. **A)** Schematic of the experiment, including collection of individual blastomeres from 1-cell and cleavage stage embryos, whole genome amplification, DNA sequencing, alignment of DNA reads to the genome, and replication timing for single cells. From zygotes, individual haploid nuclei were isolated. Early replicating areas are expected to be overrepresented relative to late replicating areas. Blue and red colors are collapsed to a one dimension in single cell replication timing profiles below. **B**) DNA replication timing in mouse embryos on chromosome 3 for maternal (M) and paternal (P) haploid nuclei from parthenogenetic zygote (N=44), fertilized zygote (N=36), maternal parthenogenetic 2-cell (N=59), 4-cell embryos (N=76), fertilized 4-cell embryos (maternal plus paternal genomes) (N=84), and mouse ESCs on chromosome 2 from (*16*), respectively. Gene density, Lamina-associated domains from zygotes and 2-cell stage embryos (*19*), origin density from mouse embryonic stem cells (*21*), are shown. vomeronasal 2 receptor (Vmn2r) gene clusters and a long neuronal gene, Fstl5, as well as an intergenic region are indicated as yellow shaded areas. **C-E**) Quantification of zygote (C), 2 cell (D) and 4 cell stage (E) LADs in correlation with late replication timing. Lamina association observed/expected (OE) >1 indicates higher lamin association than random. **F-H**) Quantification of A/B compartment in early and late replicating regions in mouse zygotes (F), 2 cell (G) and 4 cell stage (H) embryos. Positive value indicates A compartment and negative values indicates B compartment. **I-K)** Quantification of gene density in early and late replicating regions in mouse zygotes (I), 2 cell (J) and 4 cell embryos (K). **L)** Quantification of replication percentage at long genes (over 500 kb), at intergenic regions >1Mb, and at randomly selected regions. **M-O**) Quantification of origin counts in early and late replicating regions in mouse zygotes (M), 2 cell (N) and 4 cell embryos (O). Statistical test according to Mann-Whitney test. (****p<0.0001).

Clustering analysis showed that 2-cell and 4-cell embryos showed similar replication patterns for autosomes (R=0.82), and even zygotes showed relatively high correlation with 4-cell embryos (R=0.8, **Fig. S2A**). Fertilized 4-cell cleavage stage embryos and parthenogenetic 4-cell embryos, which contain only a maternal genome, showed highly similar replication patterns, suggesting that DNA replication patterns are established on the genomes of either parental origin (R=0.94, **Fig.1B, Fig. S2B**). We also made comparisons with single cell replication timing analysis from a previous study on embryonic stem (ES) cells (*16*), and found strong correlation of mouse embryo replication timing patterns with those of single mouse ES cells (R=0.83, **Fig. 1B, Fig. S2C**), with some potential local differences (arrow in **Fig. 1B**). Furthermore, we compared aggregate single cell replication timing patterns with replication timing patterns of mouse primordial germ cells (PGC), ESCs, iPSCs and mouse embryonic fibroblasts, which had been generated from a population of cells (*17, 18*). Mouse embryos showed high correlation to primordial germ cells (r=0.91, **Fig. S2D**) and iPSCs (R=0.87, **Fig. S2E**), but lower correlation was found to more differentiated cells, including mouse embryonic myoblast (R=0.79, **Fig. S2F**) and mouse embryonic fibroblasts (R=0.79, **Fig. S2G**). Thus, mammalian embryos show patterned progression of DNA replication, most closely related to primordial germ cells and embryonic stem cells.

### Late DNA replication correlates with LADs, B compartment, early replication with high gene and origin density

Lamina Associated Domains (LADs) is the first chromatin architecture pattern established after fertilization, prior to the establishment of epigenetic patterns on histones and prior to DNA methylation(*19*). In somatic cells, LADs are known to be correlated with gene-poor late replicating regions (*20*). To determine whether embryo replication timing patterns correlated with LADs, we compared zygote LADs with zygote replication timing, as well as blastomere LADs with blastomere replication timing. Both mouse zygotes and blastomeres show visible correlation between replication timing and LADs patterns (**Fig. 1B**). Late replicating regions show greater than average (OE ratio) lamina association for both zygotes and blastomeres (**Fig.1C,D,E**). We also examined correlation between replication timing and nuclear compartmentalization. In somatic cells, late replicating regions localize to the B compartment, characterized as surrounding the nucleolus and near the nuclear lamina (*20*). In both fertilized and parthenogenetic embryos, late replicating regions associate with the B-compartment (**Fig. 1B**). This association is highly significant from the first cell cycle (**Fig. 1F**), continuing through the 2-cell and 4-cell stages (**Fig. 1G, H**). Thus, the link between late replication and lamina association, as well as the segregation of replication timing according to compartmentalization into A and B compartments, begins prior to embryonic genome activation, wich occurs at the 2-cell stage in mice.

To further understand the properties of replication timing in the mammalian embryo, we compared replication timing to gene density and to origin density in ESCs. Late-replicating regions in mouse embryos, such as the gene Fstl5, a gene spanning over 600kb and expressed in the nervous system, showed low gene density (**Fig. 1B**). We quantified gene density in 100kb bins throughout the genome and found that early-replicating regions identified in zygotes and cleavage stage embryos contained significantly more protein-coding genes than late-replicating regions (**Fig. 1I,J,K**). Long genes with transcripts over 500kb and in particular intergenic regions over 1Mb were late-replicating (**Fig. 1L**, **Fig. 1B**). Regions containing long genes are intrinsically gene-poor, contributing to the low gene density of late replicating regions. However, some gene-rich regions were late replicating, in particular, gene clusters expressed in neuronal cell types: Regions encoding olfactory receptor (OR) genes and vomeronasal 2 receptor (Vmn2r) are gene-rich, but late replicating from the first cell cycle (**Fig. 1B**, **Fig. S3A-C, Table S1**). We also compared DNA replication timing in embryos to origin density, using replication origin data from mouse embryonic stem cells. We used data from mouse ES cells as these are the most closely related cell types with available origin mapping data (*21*), and as their DNA replication timing patterns correlate closely with mouse preimplantation embryos. Late-replicating regions showed significantly lower origin density than early-replicating regions in mouse embryos (**Fig. 1B**, **Fig. 1M,N,O**). This pattern also applied to gene rich regions that were late replicating: OR and Vmn2r gene clusters are origin poor in mouse embryonic stem cells (**Fig, S3D**). In contrast, the Hox gene cluster was early replicating and origin-rich (**Fig. S3C,D**). Thus, late replication correlates primarily with low origin density.

### Spontaneous chromosome break sites map to late-replicating regions in bovine embryos

To determine whether patterned DNA replication timing is also observed in other mammalian species, we chose the bovine model. We analyzed 114 bovine blastomeres harvested at the 2-7 cell stage in vitro fertilized embryos from a published dataset (*22*), corresponding to a stage immediately before embryonic genome activation in bovine (*23*) (**Fig. 2A**). As in mice, bovine preimplantation embryos showed early and late replicating regions, correlating largely with gene density (**Fig. 2B**). Yet again, the cadherin CHD6-CHD18 gene cluster region, which is expressed in differentiated neurons, was late replicating in bovine embryos (**Fig. 2A**).

**Figure 2.**
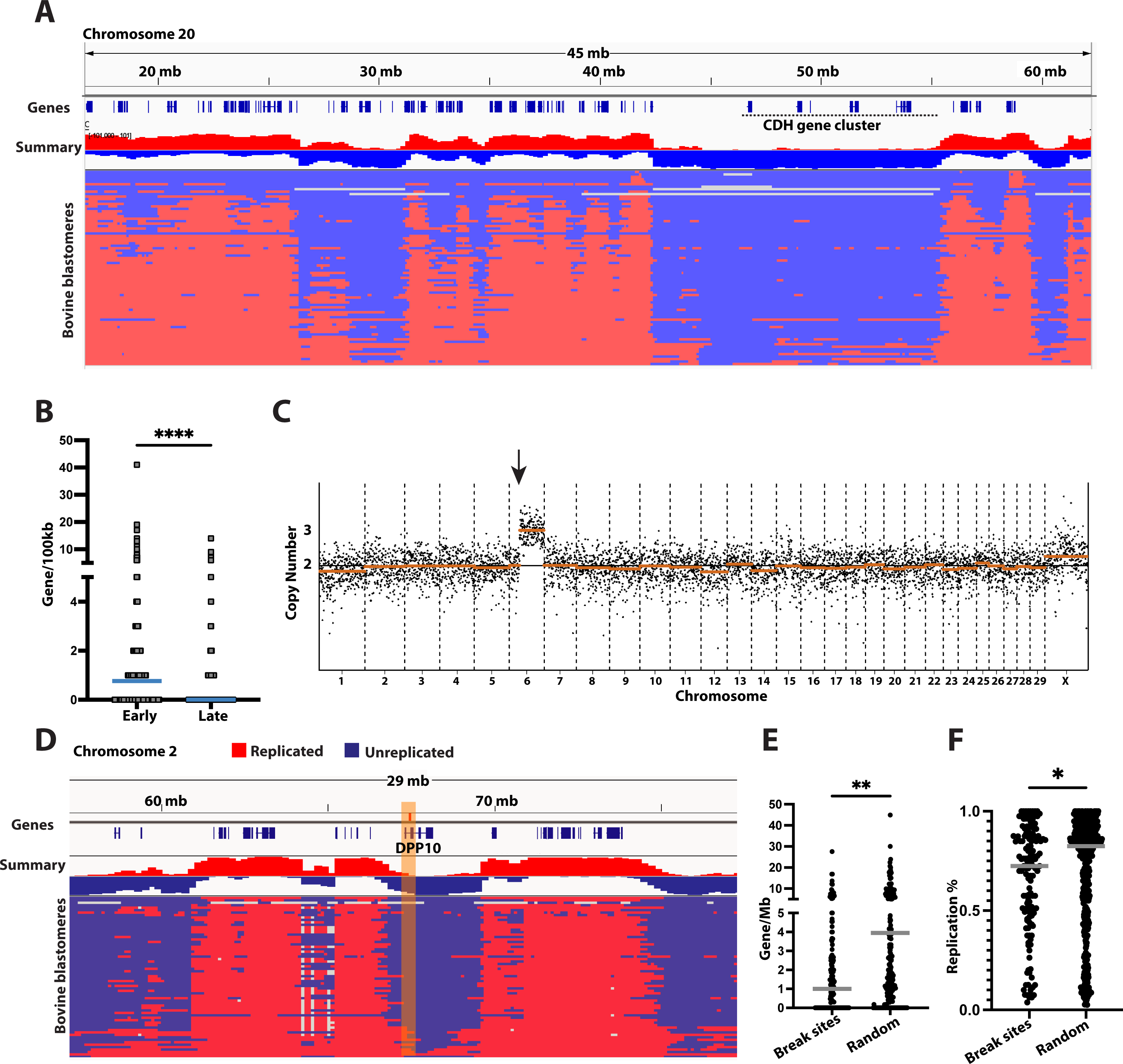
Sites of spontaneous chromosomal breakage in bovine embryos preferentially locate to late replicating regions. (**A**) Replication timing of bovine embryos on chromosome 20. Late replicating cadherin (CHD) gene cluster is indicated. **B**) Gene density of early versus late replicating regions. **C**) Karyotype of one blastomere of a day 2 bovine embryo. **D)** DNA replication timing profile in bovine embryos on chromosome 2. Top plot shows the sum of the replication status of all cells, together with gene density. The yellow bar highlights a spontaneous chromosomal break site as reported in the long fragile site gene DPP10. **E)** Quantification of gene density at spontaneous break sites and randomly selected regions. **F)** Quantification of the percentage of DNA replicated in all cells at 141 spontaneous break sites identified in bovine cleavage stage embryos and at randomly selected regions. Statistical test according to Mann-Whitney test. (****p<0.0001, **p< 0.01, *p<0.05).

In somatic cells, late-replicating, gene-poor and origin-poor regions are prone to chromosome fragility (*8, 24, 25*). Though spontaneous chromosome breakage is rather uncommon in somatic cells, replication fork slowing using low concentrations of aphidicolin is sufficient to induce frequent breakage in these regions (*26*). DNA replication fork progression in mammalian preimplantation embryos is slow even without added aphidicolin (*1, 9*), and breakage occurs spontaneously. Bovine embryos reproduce the frequent spontaneous mitotic aneuploidies and chromosome breakage seen in human (*5, 22*), but chromosomal coordinates had not previously been reported. To determine whether late-replicating regions are prone to fragility, we identified break sites in blastomeres of bovine cleavage stage embryos (**Fig. 2C, Table S2)**. We used sites of copy number transition resulting from mosaic segmental chromosome aneuploidies as a readout of the sites of chromosome breakage occurring after fertilization. We identified 141 break sites, which were located preferentially, though not exclusively to gene-poor regions (**Fig. 2D,E, Table S2**), as previously reported for human embryos (*1*). A break site in the gene DPP10 shown in **Fig. 2C**, is in direct concordance with a break site found in human embryos (*27*). Importantly, and novel in this context, bovine fragile sites show delayed replication relative to random sites (**Fig.2F**). This suggests that late replicating regions are prone to fragility in the bovine embryo.

### Paucity of dormant origins limit adaptation to replication stress at the 1-cell stage

Unlike the frequent aneuploidies and chromosome breakage in bovine embryos, no segmental chromosomal changes were found in any of the 84 mouse blastomeres, and only three (3.5%) blastomeres were aneuploid. The low frequency of spontaneous breakage makes mouse embryos amenable to studies on experimentally induced stress. We used parthenogenetic embryos with only maternal genomes, as these allow synchronous timing of cell cycle progression, and precise knowledge of the timing of activation.

In somatic cells, treatment with low concentrations of aphidicolin results in chromosome fragility due to limiting origin density in gene-poor regions of the genome (*24*). We examined origin density and replication fork speed using DNA fiber analysis in mouse embryos from the 1-cell stage to the blastocyst stage (**Fig. 3A**). DNA fibers were stained for IdU and for ssDNA to evaluate the continuity of the DNA fiber and origins were identified through evaluation of divergent sister forks (**Fig. 3B**). Median inter-origin distance was 34kb in 1-cell embryos and increased to 81kb at the blastocyst stage (**Fig. 3C**). DNA replication fork speed was evaluated by staining for both IdU and CldU (**Fig. S4A**). Origin density correlates inversely with replication fork speed through preimplantation development as described previously (*9*). Interestingly, replication fork speed was slowest at the 1-cell stage (**Fig. 3D**), suggesting that the mechanisms causing fork slowing, which are currently not known, are most active during the first cell cycle.

**Figure 3.**
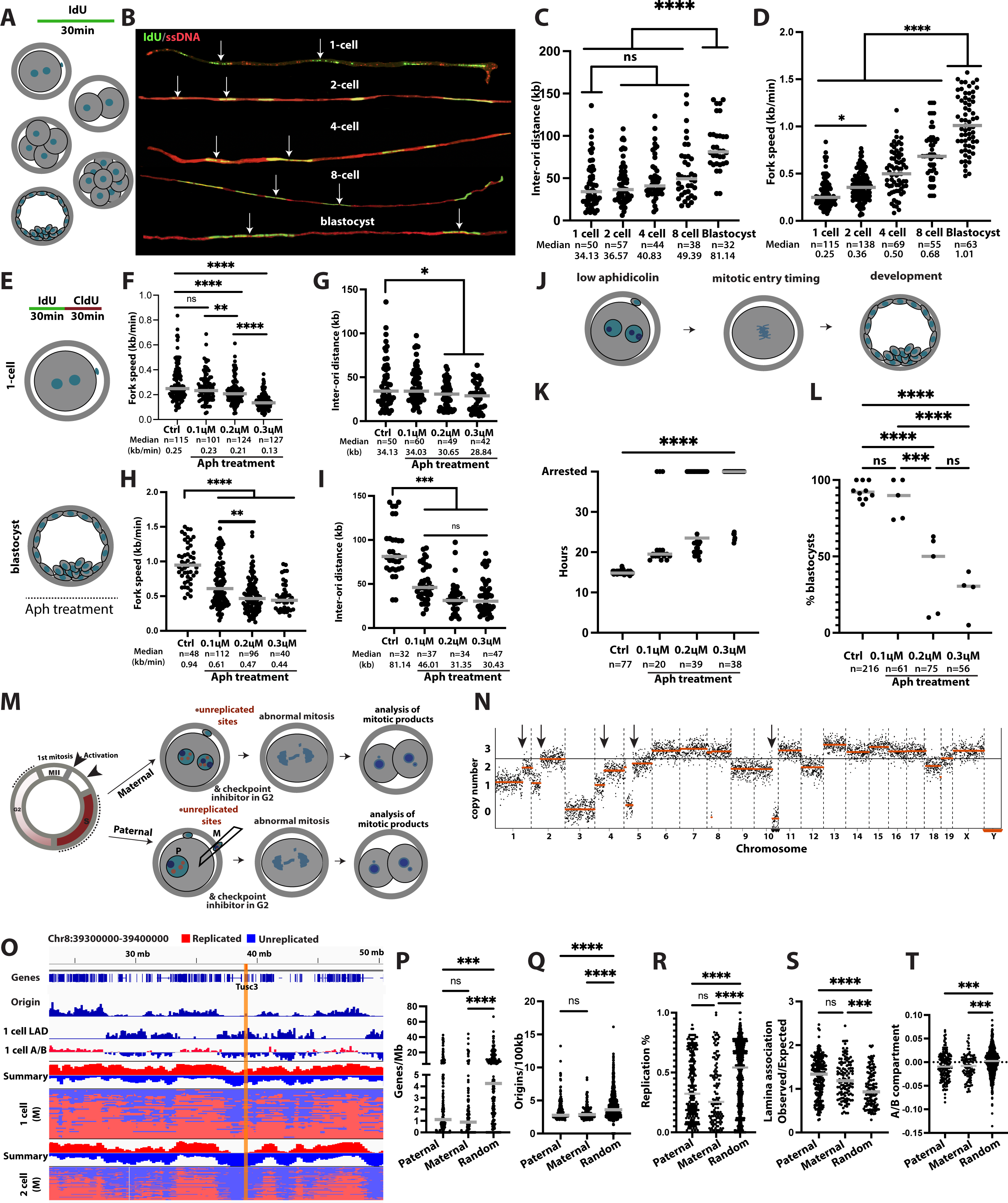
Early embryos activate few dormant origins under replication stress, affecting integrity of late replicating LADs. **A)** Schematic of the experiment. IdU is applied for 30 minutes to evaluate DNA replication fork progression and origin density at different stages. **B**) Representative DNA fibers stained for IdU and ssDNA. Arrows indicate neighboring origins. Size bar = 20kb, calculated as 2.59 ± 0.24 kbp/μm according to (*41*). **C**) Quantification of inter-origin distance depending on developmental stage. **D**) Quantification of DNA replication fork speed depending on developmental stage. **E**) Schematic of the experiment. Aphidicolin is added and embryos are labeled with sequential pulses of IdU and CldU to measure replication fork speed. **F**) Quantification of DNA replication fork speed in controls and at 0.1µM, 0.2µM and 0.3µM aphidicolin at the 1-cell stage. **G**) Quantification of inter-origin distance at indicated conditions at 1-cell stage. **H)** Quantification of DNA replication fork speed in controls and at 0.1µM, 0.2µM and 0.3µM aphidicolin at the blastocyst stage**. I)** Quantification of inter-origin distance at indicated conditions at the blastocyst stage. **J)** Schematic of the experiment. Parthenogenetic mouse embryos are exposed to low concentrations of aphidicolin and cell cycle progression is measured (**K**), as well as development after release from the drug (**L**). **K**) Timing of mitotic entry in the presence of indicated concentrations of aphidicolin. **L**) Development of mouse parthenotes after replication with indicated concentrations of aphidicolin during the first cell cycle. **M**) Schematic of the assay. Mouse embryos are incubated in low concentrations of aphidicolin throughout S-phase, resulting in unreplicated sites. For androgenetic embryos, maternal nucleus were removed 7-9 hours post fertilization. G2 checkpoint inhibition is applied at 12-14h post exposure to allow mitotic entry, resulting in an abnormal mitosis and daughter cells with an abnormal karyotype. Segmental breakpoints are a readout for unreplicated DNA at entry into mitosis. **N**) Karyotype of mouse blastomeres exposed to low concentrations of aphidicolin at the 1-cell stage on day 2 of development. **O**) Replication timing, origin density and gene density at a break site (yellow vertical bar) induced by low concentrations of aphidicolin. **P-T**) replication stress-induced break sites show lower gene density (**P**), lower origin density (**Q**), late replication timing (**R**), higher lamina association (**S**) and B compartment association (**T**) than randomly selected sites. Statistical test according to Mann-Whitney test. (****p<0.0001, ***p<0.001, **p< 0.01, *p<0.05).

To evaluate the ability of preimplantation embryos to activate dormant origins, we incubated 1-cell embryos in low concentrations of aphidicolin during the first S-phase and determined DNA replication fork speed, origin and fork density, mitotic entry timing, and developmental potential (**Fig. 3E**). Aphidicolin at the 1-cell stage reduced replication fork speed in a concentration-dependent manner from 0.25kb/min in controls to 0.21kb/min in 0.2μM and to 0.13kb/min in 0.3μM aphidicolin (**Fig. 3F**). Inter-origin density decreased only marginally from 34.13 kb to 30.27kb at the zygote stage at 0.2μM aphidicolin (**Fig. 3G)**. 0.1μM aphidicolin caused only minor, though not statistically significant changes in fork speed, origin density and fork density. Thus, the ability to activate dormant origins at the zygote stage is low, possibly because most available origins are already active.

In contrast to the 1-cell stage, aphidicolin incubation at the blastocyst stage reduced replication fork speed from ∼1kb/min to ∼0.5kb/min, causing significant changes even at low concentrations of aphidicolin (**Fig. 3H**). Origin density increased from ∼80kb to 32kb origin to origin distance, to a density observed at the 1-cell stage (**Fig. 3I**). This dramatic increase in replication origin density at low concentrations of aphidicolin shows a greater ability in blastocyst embryos to compensate for a reduction in fork speed and activate dormant origins than in 1-cell embryos, possibly because these origins are already physiologically active at the 1-cell stage.

Low replication fork speed at the 1-cell stage and a low ability to activate dormant origins may be limiting to DNA replication completion. To test this, we incubated mouse 1-cell embryos in low concentrations of aphidicolin (**Fig. 3J**), and monitored cell cycle progression. 0.1µM aphidicolin delayed entry into the first mitosis by 5 hours (**Fig. 3K**). Thus, even minor reductions in replication fork speed have a significant effect on the kinetics of cell cycle progression. At 0.2 and 0.3µM, mitotic entry was further delayed and most zygotes failed to progress to mitosis (**Fig. 3K**). Though replication fork speed was reduced by only ∼0.04kb/min at 0.2µM aphidicolin, most 1-cell embryos were arrested. Thus, mouse 1-cell embryos are highly sensitive to slowing of DNA replication fork speed below the already physiologically slow fork progression. Replication fork slowing due to aphidicolin exposure was damaging to developmental potential: when embryos were released from aphidicolin after 24 hours of exposure, most 1-cell embryos released from 0.2µM or 0.3µM aphidicolin failed to develop to the blastocyst stage (**Fig. 3L**).

### LADs and genomic regions in B-compartment are sensitive to replication fork slowing from the first cell cycle

To identify the genomic regions with limiting origin density in the early embryo, we used low concentrations of aphidicolin throughout the first S-phase and added G2 checkpoint inhibitors to facilitate mitotic entry. Upon entry into mitosis, unreplicated sites result in chromosome breakage and aneuploidy in cleavage products, which can be determined through copy number analysis (**Fig. 3M**). We used parthenogenetic embryos as well as androgenetic embryos for these studies to determine whether potential patterns are observed on both maternal and paternal genomes. Low concentrations of aphidicolin resulted in abnormal chromosome content with varying copy number of each chromosome (**Fig. 3N**). G2 checkpoint inhibition alone results in aneuploidy in only a third of all cells (*1*), and thus most chromosomal abnormalities in this assay are caused by the slowing of replication fork progression by aphidicolin during S-phase. Segmental errors manifest as copy number transitions within a chromosome, thereby providing the coordinates for breakage (**Fig. 3N, Table S3**). For instance, **Fig. 3O** shows the coordinates for a copy number transition at the gene Tusc3 together with the corresponding DNA replication timing profile in embryos and origin density. Coordinates for a total of 125 copy number changes were identified and gene density and origin density quantified (**Table S3**). Sites of chromosome breakage were generally gene-poor (**Fig. 3P**) or within an OR gene cluster (**Fig. S3B**), origin-poor (**Fig. 3Q**), and preferentially located to late-replicating regions (**Fig. 3R**). Furthermore, chromosome breakages were associated with LADs (**Fig. 3S**) and were enriched in the B compartment (**Fig. 3T**).

Taken together, these results show that there is a limited ability to respond to replication stress at the 1-cell stage, which is most consequential in late replicating regions that are origin-poor, gene-poor, and associated with the nuclear lamina, predisposing these regions to breakage. This pattern is established already in the first cell cycle, on the maternal as well as the paternal genome. Importantly, this pattern is congruent with patterns of spontaneous chromosome breakage in cow and human embryos, and thus physiologically relevant.

### Replication gaps locate to late replicating, gene poor, origin poor regions, associated with the nuclear lamina and the B compartment from the first cell cycle

DNA replication stress results in a requirement for G2 and mitotic DNA synthesis in somatic cells (*28*). Spontaneous replication stress in the embryo is associated with DNA synthesis in G2 phase, involving gaps of ∼1kb or less (*1*). These are likely sites of postreplicative repair.

To evaluate the cytological location of postreplicative DNA synthesis and repair, we stained fertilized mouse zygotes in late G2 phase, immediately prior to entry into mitosis for *γ*H2AX, and measured proximity to the nuclear envelope and the nucleolus. Foci localized to the nuclear lamina and the nucleolus (**Fig. 4A)**. Only 10% of all foci (n=331) localized within the A compartment, away from either the nuclear envelope or the nucleolus. Of note, a greater portion of foci is associated with the nucleolus (53%) than with the nuclear lamina (37%) (**Fig. 4B**), adding up to 90% B-compartment (**Fig. 4C**). This pattern was apparent on both maternal and paternal genomes, as identified by ∼1μm greater diameter. Thus, cytological patterns of delayed replication predisposing to fragility are established during the first cell cycle, in accordance with regions of late DNA synthesis, which occurs near the nuclear lamina and around the nucleolus (*29*).

**Figure 4.**
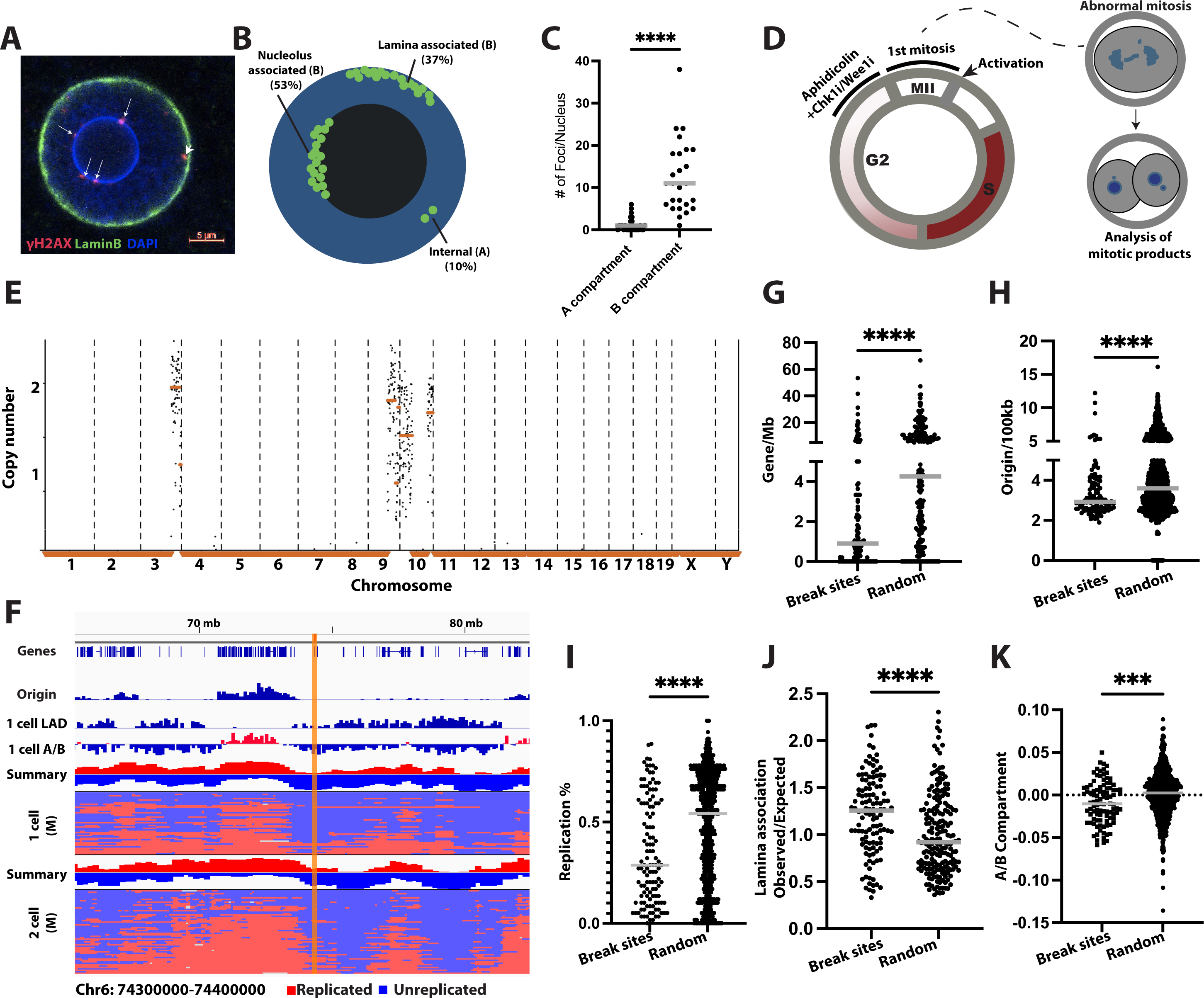
Origin poor, lamina associated regions show incomplete replication in late G2 phase and are prone to fragility. **A**) Immunostaining of a fertilized nucleus at late G2 phase for *γ*H2AX and lamin b. Arrow points to foci localized on nucleolus and arrow head points to foci localized on nuclear envelope. Size bar = 5µm. **B**) Schematic of foci location. **C**) Quantification of foci (n=331) in different compartments in 35 nuclei; on and in proximity to nucleolus and nuclear envelope (B compartment), and A compartment. **D)** Schematic of the experiment. Aphidicolin is added at high concentration at the end of the first cell cycle to inhibit DNA synthesis in late G2 phase to probe for regions with incomplete replication at that time point. CHK1 inhibition or WEE1 inhibition facilitates mitotic entry despite unreplicated DNA, resulting in chromosome breakage, and mitotic products with micronucleation and aneuploidy. Coordinates of chromosome breakage are determined using low pass single cell genome sequencing, providing a readout for the sites of incomplete replication. **E**) Chromosomal content analysis of a micronucleus isolated from a mouse blastomere after transition through the first mitosis. **F**) DNA replication timing profile in mouse embryos on chromosome 6. Top shows the sum of replication status of all cells, together with gene and origin density, LADs and A/B compartment. The yellow bar highlights the location of a chromosomal break site caused by inhibition with aphidicolin in G2 phase. (**G-K**) Sites of G2 replication identified through mapping of break points in mice show lower gene density (**G**), lower origin density (**H**), later replication timing (**I**), higher lamina association (**J**) and B compartment association (**K**) than randomly selected sites.

To determine the location of the sites requiring G2 DNA synthesis after an unperturbed first S-phase in mouse embryos, we interfered with DNA replication completion in G2 phase using high concentrations of aphidicolin combined with G2 checkpoint inhibition (**Fig. 4D**). Sites that remain unreplicated incur chromosome breakage in mitosis, resulting in micronucleation and aneuploidy. Genomic coordinates of segmental copy number changes caused by chromosome breakage at unreplicated sites, provide a readout for the location of unreplicated DNA in G2 phase. We used parthenogenetic mouse 1-cell embryos containing only maternal genomes for these studies, as both paternal and maternal genomes cytologically show the same patterns of postreplicative repair, and as partenotes allow precise knowledge of the timing after activation.

Blastomeres of the resulting 2-cell embryos showed frequent chromosomal aneuploidies with segmental errors (**Fig. 4E, Table S4**). **Fig. 4F** shows a break site in a region that is both origin-poor and gene-poor, as well as late replicating. In aggregate, sites of chromosome breakage were found to be gene-poor (**Fig. 4G**), origin-poor (**Fig. 4H**), and also replicated later than randomly selected sites at the cleavage stage (**Fig.4I**). In addition, breakage occurred at neuronal gene clusters, including olfactory receptor gene clusters (OR) and vomeronasal receptor gene cluster (Vmn2r) (**Table S4**). We also analyzed the correlation of break sites with LADs, showing higher association with zygote LADs in chromosome breakages compared with random sites (**Fig.4J**). Furthermore, break sites preferentially associated with the B compartment (**Fig. 4K**).

Chromosomal breakages resulting from replication fork slowing (**Fig. 3**) or resulting from incomplete replication in G2 phase (**Fig. 4**) showed concordance, affecting areas with late replication timing, low origin density, low gene density, nuclear lamina and B compartment association. Sites of direct concordance were found at Prkg1, Lrp1b, A1cf and at two noncoding regions (**Table S5**). There is also direct concordance with spontaneous fragility in other species: for example, chromosome breakage at Lrp1b was found in untreated fertilized human embryos (**Table S5**) (1). Thus, late replicating, origin-poor, lamina-associated regions of the genome are prone to incomplete replication and to chromosome breakage. This pattern is apparent already in the first cell cycle.

## Discussion

Here we show that DNA replication in early mammalian embryos progresses in a defined pattern with early replicating gene-rich regions and late-replicating, origin-poor regions containing long neuronal genes. In addition to long intergenic regions, gene clusters expressed in highly specialized neuronal cell types, including olfactory receptor genes, and vomeronasal receptor genes and cadherin genes were also late replicating. Replication timing patterns in mice showed high correlation with mouse embryonic stem cells, and lower correlation with differentiated somatic cells. Furthermore, we show that late replicating regions have association with the nuclear lamina and preferentially associate with the B compartment. Therefore, DNA replication timing in early mammalian embryos follows basic principles applicable to more differentiated cell types.

The early embryo provides a unique model system to study the establishment of epigenetic regulation of the genome. Previous studies have shown that lamina associated domains are becoming apparent from the 1-cell stage (*19*), while DNA methylation and histone patterns are established only later in development. Similarly, compartments of accessible and inaccessible chromatin are beginning to be established as early as the 1-cell stage (*30*), as are patterns of chromatin architecture (*31*). However, these studies also found that the patterns at the 1-cell stage show lower segregation than at the cleavage stage. We find that this is also reflected in the DNA replication patterns: single nuclei isolated from fertilized zygotes show late and early replicating regions concordant with those of cleavage stage embryos, even as the amplitudes of these differences are lower.

The maternal genome has been reported to show delayed lamina association compared to the paternal genome at the 1-cell stage, gradually assimilating to the paternal genome after the 8-cell stage (*19*). While differences between paternal and maternal genomes are possible, basic principles determining DNA replication patterns are conserved between the two genomes: cleavage stage embryos, fertilized or parthenogenetic, show similar patterns of DNA replication timing, both correlating with LADs. Cytological analysis is in concordance with these findings: from the 1-cell stage, both paternal and maternal genomes show organized spatial and temporal progression of DNA replication, with late replicating regions associated with the nucleolus and the nuclear lamina (*29*). Adding to that, we show that both DNA damage foci in late G2 phase, as well as very late replicating regions prone to breakage in mitosis, show a distinct pattern of nuclear lamina and nucleolar association, including in embryos with only the maternal genome.

Studies on nuclear lamina association using Lamin-DamID may underestimate the compartmentalization of the genome in the early embryo, because of a very prominent nucleolus. Recent studies in cultured cells show that domains with nucleolar association also show late DNA replication timing (*32*). Furthermore, the quality of the data may be more variable, than in cells that can be obtained in greater abundance.

The question of developmental timing when DNA replicaiton patterns are established is relevant to the question of cause and consequence. Taken together, the data presented here show that the segregation of early and late replicating DNA are among the earliest epigenetic features installed on the mammalian genome during development, and that both the nuclear lamina and the nucleolus are involved in establishing these patterns.

In addition to mice, we chose bovine embryos for analysis because the genome of bovine embryos is very unstable during cleavage development. Remarkably, bovine embryos reproduce the spontaneous pattern of breakage seen in human embryos, occurring preferentially in gene-poor regions. Breakages at concordant sites between bovine and human embryos were found on KCNMA1 and FHOD3 locus, which are both involved in nervous system development (*33, 34*), in FHIT, a fragile site found in somatic cells exposed to aphidicolin and in tumor cells (*26, 35*) and at DPP10, a long neuronal gene, and a hotspot of copy number variants (*36*). Fragile sites in bovine embryos preferentially, though not exclusively, locate to late-replicating regions.

In early mammalian embryos, the replication program is strained from intrinsic replication stress and has a limited ability to activate additional origins in response to exogenous stress. Regions in the genome most prone to breakage show a relative paucity of origins and continue to undergo DNA synthesis in G2 phase. These patterns of G2 replication and fragility are established from the first cell cycle, in association with the nuclear lamina and the nucleolus. Lamina associated regions are known to be fragile in the germ line and in cancer cells. For instance, the genes DMD and PARK2 are lamina associated regions with structural changes observed in the germ line in patients, and in tumor cells (*37*). Breakage at both of these loci has been observed in mammalian embryos (this study & (*1*)). Our observations raise the question whether clonal genetic change in the soma and the germ line arises after fertilization as a result of replication stress as early as the 1-cell stage. Patterns of DNA replication and the resulting patterns of fragility in totipotent cells may shape mammalian genome evolution in a manner less available to lower vertebrates, as these establish replication patterns only after hundreds of nuclei have been formed (*12*). Of note, mutation rates are higher in late replicating regions than in early replicating regions (*18*), and thus these higher mutation rates would occur from the first cell cycle. Our finding that numerous genes expressed in the nervous system replicate late may be particularly relevant for generating genetic and phenotypic diversity in the brain. Future studies should examine the causal relationships of fragility patterns, DNA replication timing, germline mutations, and other layers of epigenome regulation in the early mammalian embryo.

## Materials and Methods

### Mouse embryos

Mouse oocytes were obtained from B6D2F1 females 5-8 weeks of age from Jackson laboratories (stock # 100006), after hormonal stimulation of 5 IU PMSG per mouse, followed 48h later by 5 IU hCG per mouse. Oocytes were dissected from oviducts 14-16h post hCG application and dissociated in hyaluronidase. In vitro fertilization (IVF) was performed using sperm from 21-20 week old male mice extracted from the epididymis, in Global total for fertilization (LGTF-050). Artificial activation of mouse oocytes was performed using 1µM ionomycin in Global Total for 5 min. at 37 deg., followed by 3.5h 10µg/ml puromycin and 5µg/ml cytochalasinB, followed by 1.5h in cytochalasinB only. Mouse parthenotes were then cultured in Global Total (LGGT-030) at 37 deg. In 5% CO2 atmosphere and harvested 8 hours post activation/fertilization for zygote stage, 20-23 hours post activation for 2 cell stage, and 28 hours for 4 cell stage. Zygotes and blastomeres were isolated using laser-assisted dissection in PGD medium (LPGG-020). For zygotes, 274 single maternal and paternal nuclei isolated from 2PN embryos at 7-9h were analyzed and 80 passed quality controls (described below). We estimate a time window of several hours for fertilization after insemination, which could underly the difference in signal amplitude between parthenotes and fertilized embryos. 88 blastomeres from 4-cell stage fertilized embryos were sequenced, and 84 passed quality controls. In addition, 60 and 83 parthenogenetic blastomeres from 2-cell and 4-cell embryos were analyzed using sequencing and 59 and 76 passed quality controls.

Aphidicolin incubations were done from 3h after artificial activation for low concentrations of aphidicolin throughout the first cell cycle, and at 2µM and beginning at 11-12h post activation for G2 incubations.

Immunostaining was performed for LH2AX using mouse monoclonal antibody Millipore Cat# 05-636 and for Lamin B1 using mouse monoclonal antibody Proteintech Cat# 66095-1-Ig. Distance of foci to the nuclear envelope or the nucleolus was measured using Zeiss Zen program.

All animal research has been reviewed and was approved by the Columbia IACUC.

### Bovine embryos

Bovine blastomere data were downloaded from Sequence read archive (PRJNA577965 (*22*)). Of 114 bovine blastomeres, 69 passed quality controls, all of which were cells harvested at the 2-7 cell stage. These data were used for DNA replication timing analysis and for the identification of 110 breakpoints (**Table S2**). In addition, for break point identification, bovine oocytes were obtained from DeSoto Biosciences as maturing oocytes retrieved from ovaries, and shipped in maturation medium consisting of TCM199, FSH and 10% FBS. At 37h post maturation, bovine oocytes were activated using 5μM ionomycin for 5 minutes, followed by 5µg/ml CytochalasinB plus 6-DMAP for 4.5h. Culture was performed at 38.5deg. at 5% CO2 in KSOMaa (MR-106-D, EMD Millipore) with 5% FCS. Parthenogenetic blastomeres were harvested at the 8-16 cell stage for the mapping of 31 breakpoints (**Table S2**).

### Single cell sequencing and library construction

Individual cells or nuclei were collected on the heated stage @ 37deg.C (Tokai Hit) of an Olympus IX71 inverted microscope equipped with Narishige micromanipulators and a zona pellucida laser (Hamilton-Thorne). Single nuclei were isolated by lysis and dissection of different nuclei using two 20μm diameter micropipettes (Origio). Single embryo cells or nuclei were placed manually in single wells of a 96-well plate containing 9 μl of lysis buffer, prepared as a master mix of 798μl H2O, 6 μl of 10 mg/mL Proteinase K solution (P4850, Sigma-Aldrich), and 96μl 10X Single Cell Lysis and Fragmentation buffer (L1043, Sigma-Aldrich). Single cells / single nuclei were lysed by heating 96-well plates containing single cells/single nuclei at 50°C for 1 hour, followed by incubation at 99°C for four minutes using a PCR thermocycler. A degenerate oligo-nucleotide primed PCR (DOP-PCR) amplification protocol that allows inline indexing of WGA DNA was applied (15, 31). WGA DNA was subsequently processed for Illumina library sequencing preparation using TruSeq indexing of the NEBNext Ultra II DNA Library Prep Kit for Illumina (New England Biolabs). NEBNext Multiplex Oligos for Illumina (96 index primers) were used for four amplification cycles. Quality control for the library was done using Qubit dsDNA HS Assay Kit (Invitrogen, Carlsbad, CA, USA) and library size distribution was confirmed using a 2100 Bioanalyzer DNA 1000 Kit (Agilent, Santa Clara, CA, USA). A unimodal curve centered around 300 – 500 bp was scored as a successful library preparation. Subsequently, 30 μL of each pooled library was sent for sequencing at a concentration of 20ng/μl. DNA sequencing was performed by Genewiz using Illumina HiSeq 4000, 2x150bp, targeting a coverage of ∼1 million reads per cell/nuclei.

### Single cell copy-number inference and break sites annotation

Sequencing data were demultiplexed according to unique barcodes and subsequently aligned to the mouse/bovine genome build (mm10/bosTau8). Analysis of DNA sequencing data was performed as previously described (*1*). Briefly, the genome was partitioned into 500kb bins and the mapped reads were sorted, indexed and counted within the genomic bins. Additionally, copy number analysis was performed using R package QDNAseq v1.26.0 (*38*) to partition the genome into 50kb and 100kb bins. Break points were annotated at the transition points in segmented read count data. Only reciprocal break sites that were found in concordance between the two segment calling methods were included in downstream analysis.

### Replication timing analysis, gene density, origin density and LADs

The replication timing method was adopted from the scRepli-seq (*39*). Briefly, the sequencing reads were aligned to the reference genome (mm10/bosTau8). MII phase oocytes were used as reference to even copy numbers across the genome. In bovine blastomeres, G1 cells were called according to (*39*) from the dataset PRJNA577965 (*22*). The mapped reads were sorted, indexed and RT scores were calculated by the control G1 cells (GSE108556). Briefly, median of the ‘absolute deviations from the median’ (MAD) scores were used as quality control to ensure read number variability was low (<0.3) for G1 phase cells and moderate (∼0.4-0.8) for S phase cells. The replication timing data were binarized as replicated versus un-replicated regions based on the replication percentage. The replication percentage was defined as the percentage of samples replicated at each 100kb bin. Early replicating regions were defined as >50% replicated and late replicating regions were <50% replicated.

Gene density comparison was performed by randomly selecting locations within mm10/bosTau8 genome assembly using bedtools v2.30.0 (*40*). Complete transcripts and adjacent intergenic areas were included to calculate the density of protein-coding genes.

Mouse ESC origin data were obtained from (GSE68347 (*21*)). Replication timing of mouse ESCs and differentiated cells from whole genome sequencing data was inferred using TIGER (*17*). Single cell replication timing data were obtained from (*16*).

Mouse zygote and 2-cell stage embryo non-allelic LADs data were obtained from (GSE112551). For statistical analysis we used Mann-Whitney test and one-way Anova as indicated in Legends.

### DNA fiber analysis

We used activated mouse oocytes, cleavage stage as well as blastocyst stage embryos for analysis. Previous studies have found no difference in DNA replication fork speed whether fertilized or activated embryos were used (*1*). Cleavage and blastocyst stage embryos were asynchronous in their cell cycle at incubation.

For fork speed assay, the mouse embryos were incubated with 25mM IdU 30min and washed twice, and then treated with 25mM CldU for 30min. For fork density and origin to origin distance assays, cells were only exposed to 25uM ldU for 30min. Using Acidic Tyrode’s solution (MR-004-D) to digest zona pellucida in a 4-well dish at room temperature for 5-10 minutes, and then neutralized in culture medium. The cells were collected in 1-2ml medium and then placed in PCR tubes. Adding 20 ml of fresh pre-warmed (30 LJ) spreading buffer (0.5% SDS, 50 mM EDTA, 200mM Tris pH 7.4) to lysed cells, incubated for 6-8 min at RT and then stretched them on pre-cleaned microscope slides. Slides were fixed in pre-cooled methanol: acetic acid=3:1 for 2 min at RT and air dried at RT, incubated in 2.5M HCl for 50 min, and rinsed 5 times with PBS, and then blocked sides with 3% BSA in PBS for 1h. The slides were treated with anti-BrdU, anti-IdU and anti-ssDNA antibodies for 1h, and rinsed 3 times in PBS. Incubated with secondary antibodies for 1h, mounted with ProLong Gold Antifade and let dry overnight at RT. The fiber tracks were imaged on a Zeiss fluorescence microscope at 63X magnification and measured using ImageJ software v1.53. The length of each track was measured manually by ImageJ software. The pixel values were converted into μm using the scale bar generated by the microscope software. The DNA fiber length was calculated as follows: 2.59 ± 0.24 kbp/μm according to Jackson et al (*41*).

## Data availability

Mouse and bovine blastomere sequencing data are available at SRA accession number PRJNA874697.

## Author Contributions

SX, NW, and DE designed the study. SX and MZ performed library preparations, SX performed data analysis, NW performed DNA fiber analysis. SX, NW and DE performed embryology. AK assisted with data analysis, TB contributed single cell DNA amplification reagents and expertise, JG provided help with DNA fiber analysis. SX and DE wrote the manuscript with contributions from all authors.

## Acknowledgments

We thank Stepan Jerabek for critical reading of the manuscript. This work was supported in part by the National Institutes grant 1R01GM132604 to DE, and by NYSTEM grant #C32564GG. TB was supported by the William C. and Joyce C. O’Neil Charitable Trust, Memorial Sloan Kettering Single Cell Sequencing Initiative. We thank Alberto Ciccia and Angelo Taglialatela for helpful discussions and for providing microscopy resources.

**Materials and Correspondence**: requests for materials or additional information should be addressed to DE at

## Supplemental Figure Legends

**Fig. S1.**
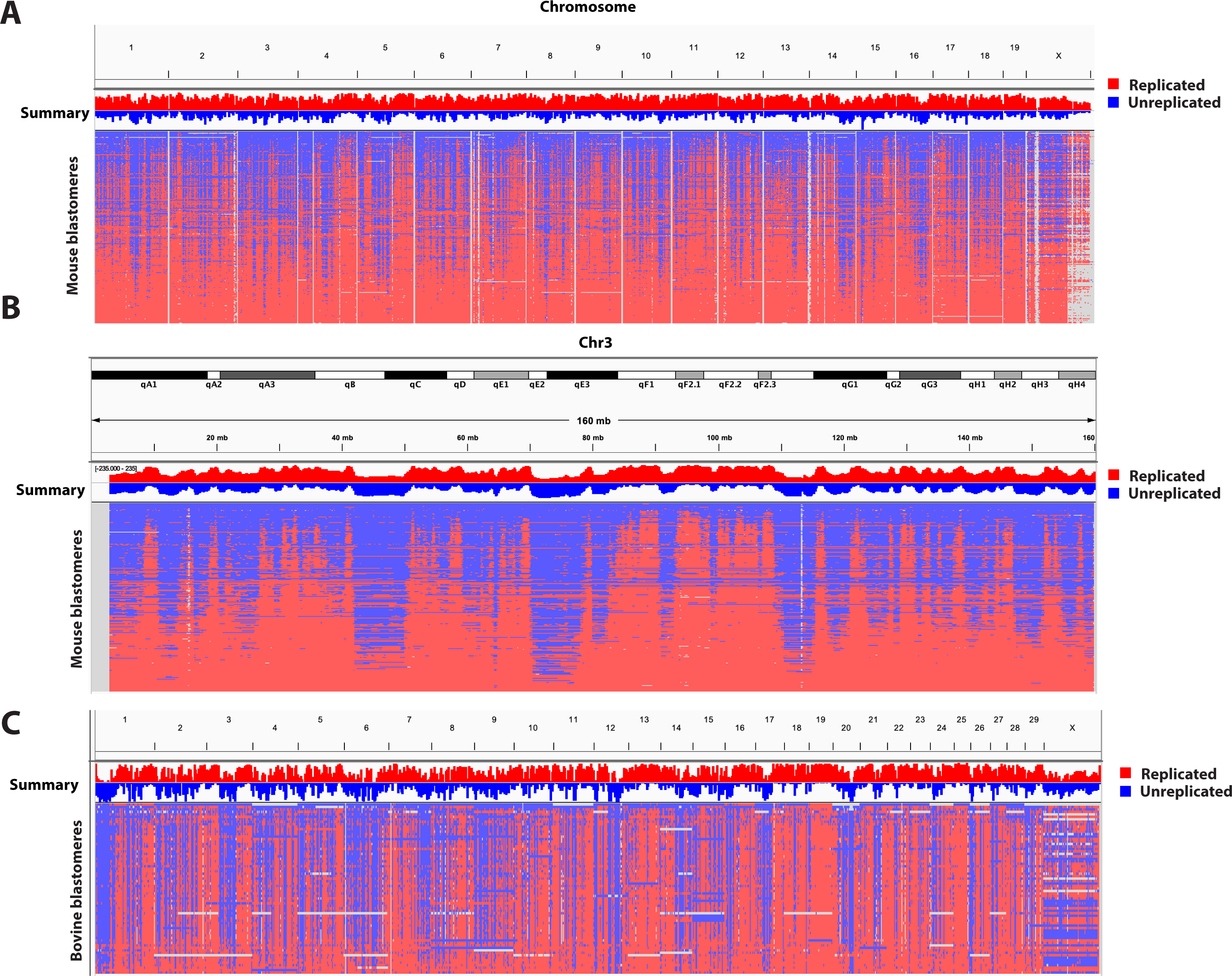
Replication timing profile of mammalian cleavage stage embryos show a temporal progression through S-phase. **A, B)** Cumulative DNA replication timing profiles of all 2-cell and 4-cell mouse embryos, maternal and paternal genomes (n=142). **(B)** Single chromosome view of the same samples shown in A. **(C)** DNA replication timing profile of all chromosomes of bovine blastomeres from day 2-3 bovine embryos (n=69). Copy number of the X chromosome is variable in mouse or bovine. Individual cells (blastomeres) are shown ordered according to the percent replicated genome. Top plot shows summary/integration of all cells as a copy number plot.

**Fig. S2.**
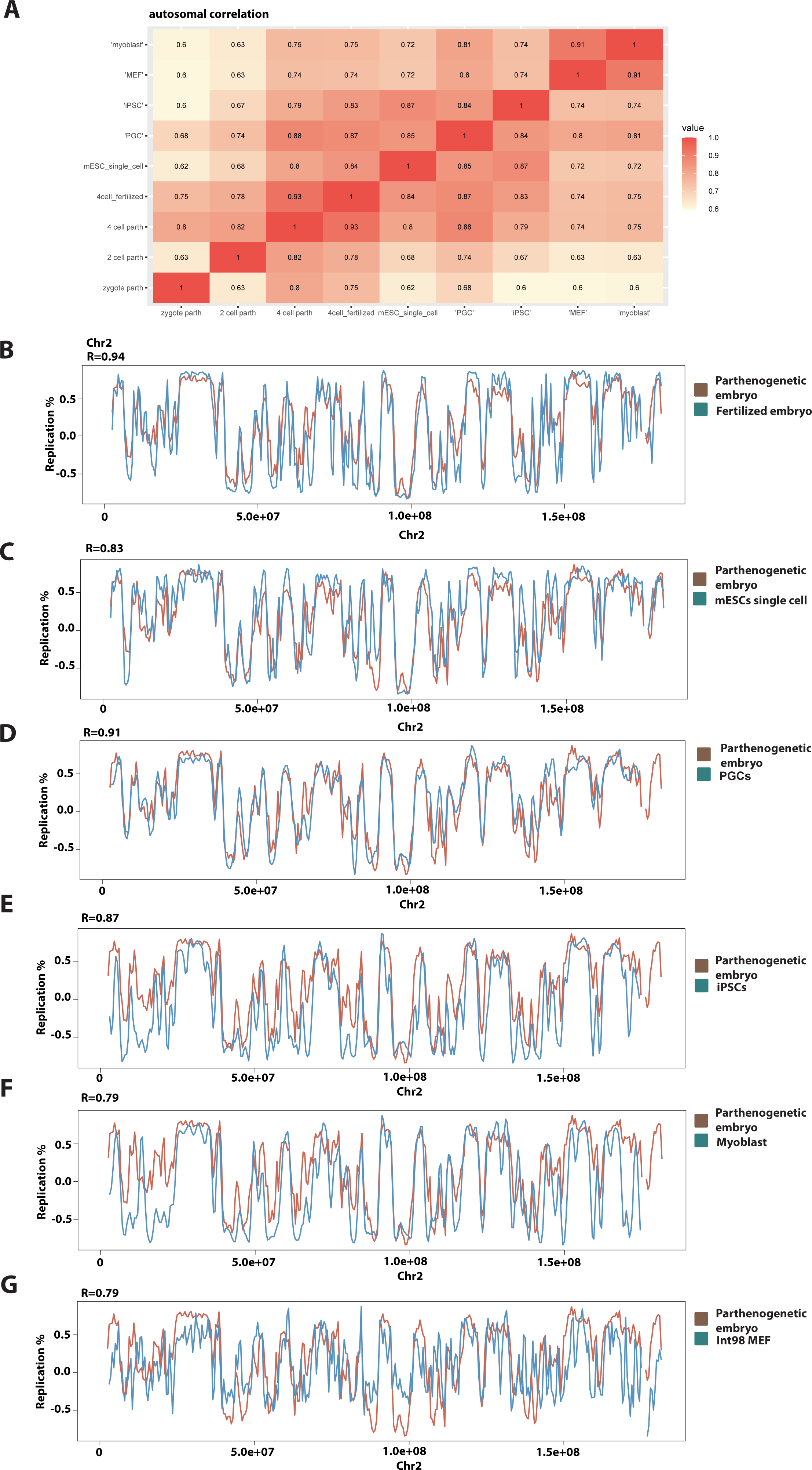
Mouse embryo replication profile has higher correlation with ES cells and primordial germ cells than with differentiated cells. A) Summary of genome replication timing profile correlation between developmental stages and cell types. Values indicate Spearman’s R. **B-G)** Replication profile correlation comparing mouse day 2 parthenogenetic embryos with **B)** fertilized mouse embryos, **C)** single cell mouse embryonic stem cells (mESCs), **D)** primordial germ cell (PGC), **E)** Mouse induced pluripotent stem cells (iPSCs), **F)** Myoblasts, and **G)** Mouse embryonic fibroblasts (MEFs). Statistical test according to Spearman correlation test.

**Fig S3.**
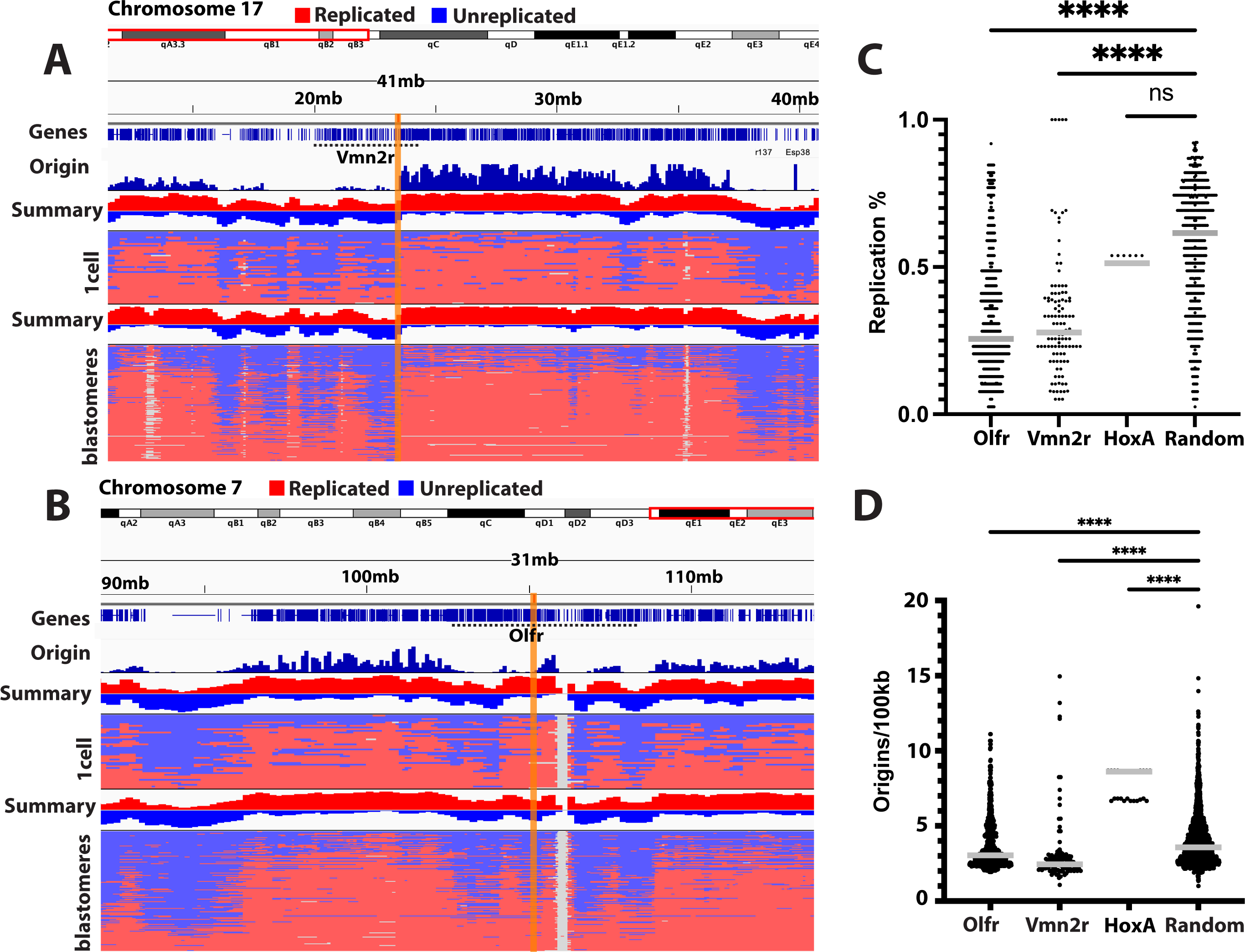
Gene clusters at vomeronasal 2 receptor (Vmn2r) and olfactory receptor (OR) genes show late replication, despite high gene density. DNA replication timing profiles, gene density and replication origin density at Vmn2r **(A)** and OR **(B)** gene clusters. The yellow bar highlights chromosomal break sites identified in mouse embryos. **C)** Quantification of the percentage of replicated DNA in all cells at gene clusters and at randomly selected regions. Lower values equate later replication timing. **D)** Quantification of replication origin density at gene clusters and randomly selected regions. (**p<0.01, ****p<0.0001). Statistical test according to Mann-Whitney test.

**Fig S4.**
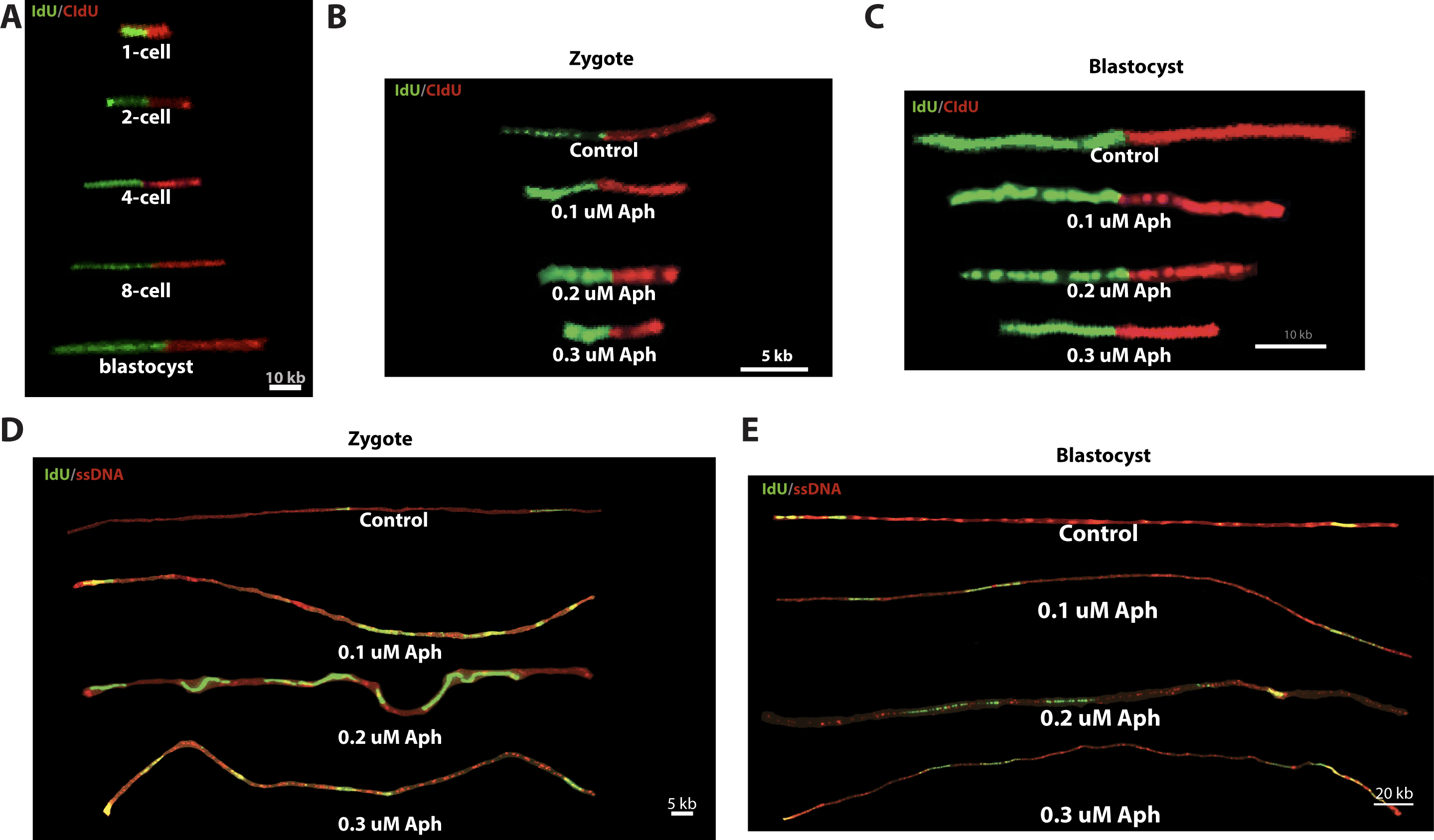
Representative images of DNA fibers from controls and at 0.1µM, 0.2µM and 0.3µM aphidicolin at the preimplantation-stage embryos. (**A-C)** DNA fibers were stained for IdU and CidU antibodies. (**D)** and (**E)** were stained for IdU and ssDNA antibodies. Developmental stages and conditions are indicated. Size bar is indicated, calculated as 2.59 ± 0.24 kbp/μm according to (*41*).

## Supplemental Tables

**Table S1. Replication timing and origin density at gene clusters, long genes over 500kb and intergenic regions over 1Mb.** (Related to Figure 1)

**Table S2. Chromosomal coordinates of spontaneous bovine break sites.** (Related to Figure 2)

Blastomeres were analyzed between the 2-cell and the 12-cell stage. Gene density calculation is based on the number of protein-coding genes.

**Table S3. Chromosomal coordinates of mouse break sites induced by low concentrations of Aphidicolin.** (Related to Figure 3)

Zygotes and Blastomeres at the 2-cell stage were treated throughout S phase with 0.2-0.4 µM of aphidicolin as indicated. Gene density calculation is based on the number of protein-coding genes.

**Table S4. Chromosomal coordinates of break sites induced through interference with G2 DNA replication in mouse embryos.** (Related to Figure 4). Blastomeres were treated with 2 µM of aphidicolin at G2 phase. Gene density calculation is based on the number of protein-coding genes.

**Table S5. Multispecies spontaneous and aphidicolin-induced break sites concordant regions.** (Related to Figures 2-4). Identified break sites were converted between species using the UCSC genome browser function to identify syntenic regions in the counterpart genome.

